# AlphaSync is an enhanced AlphaFold structure database synchronized with UniProt

**DOI:** 10.1101/2025.03.12.642845

**Authors:** Benjamin Lang, M. Madan Babu

## Abstract

Synchronizing protein structure predictions with sequence databases is crucial for their accuracy. We developed AlphaSync (alphasync.stjude.org), a pipeline and database that provides updated AlphaFold structure models for currently 39,961 UniProt proteins, including isoforms. Moreover, AlphaSync enhances 2.6 million structures with residue-level features, including solvent accessibility, dihedral angles, intrinsic disorder predictions, and 4.7 billion atom-level noncovalent contacts. AlphaSync’s user-friendly interface enables structural analyses at scale across up-to-date proteomes for 42 species.

## MAIN TEXT

AlphaFold 2 has revolutionized structural biology by providing highly accurate structural models from protein sequences^1^. Currently, 214 million proteins have predicted structures available in the AlphaFold Protein Structure Database (AFDB)^2^. However, maintaining synchronization between these structural models and available protein sequences is crucial, and it is nontrivial to detect how many sequences and structures have become outdated. For the human canonical reference proteome and UniProt-reviewed proteins^3,4^, we found that 633 proteins (3.1%) had been newly added or updated with new sequences in the UniProt database since the latest release of the AlphaFold Protein Structure Database in October 2022. These included at least 60 clinically significant genes in genetic disorders^5^ and 12 genes causally implicated in cancer^6^, such as BRCA2, MYC, AXIN2 and ATRX. Moreover, at least 52 newly added proteins are of unknown function while at least 6 are part of a primate-specific gene family, FAM90A. Beyond missing new biology, outdated sequences lead to mismatches in residue numbering and identity, errors in communication and interpretation, and inaccurate structures.

To address the important yet often overlooked problem of outdated structural models and to provide an updated reference database, we developed AlphaSync, a computational pipeline, database, API and web interface that provides updated AlphaFold structure predictions for current UniProt protein sequences (**Figure 1A**). AlphaSync uses perfect sequence matching to assign structure models in the AlphaFold Protein Structure Database to UniProt sequences and predicts new structures where needed. AlphaSync currently provides complete proteome-wide structures including isoforms for 42 species, including human, model organisms (including mouse, *D. melanogaster, C. elegans, A. thaliana, O. sativa*, and *S. cerevisiae*), and many human pathogens (**Supplementary Table 1**). To achieve full coverage of these species, 69,045 new structure predictions for 39,961 proteins were made using AlphaFold 2 on sequences from UniProt release 2025_01 (large proteins as multiple fragments) (see **Supplementary Table 2** for parameter details). The structure models are available for download from the AlphaSync website to allow easy updating of existing collections of structures. In addition, AlphaSync includes reviewed UniProt proteins, and proteins with human orthologs from 200 representative species from the Ensembl Comparative Genomics pipeline, whose sequences had matches in the AlphaFold Protein Structure Database. This unique dataset, with orthologous sequences and structure models for all human proteins, will accelerate studies on structure-function relationships and evolution of function for a wide range of protein families.

**Figure 1:**
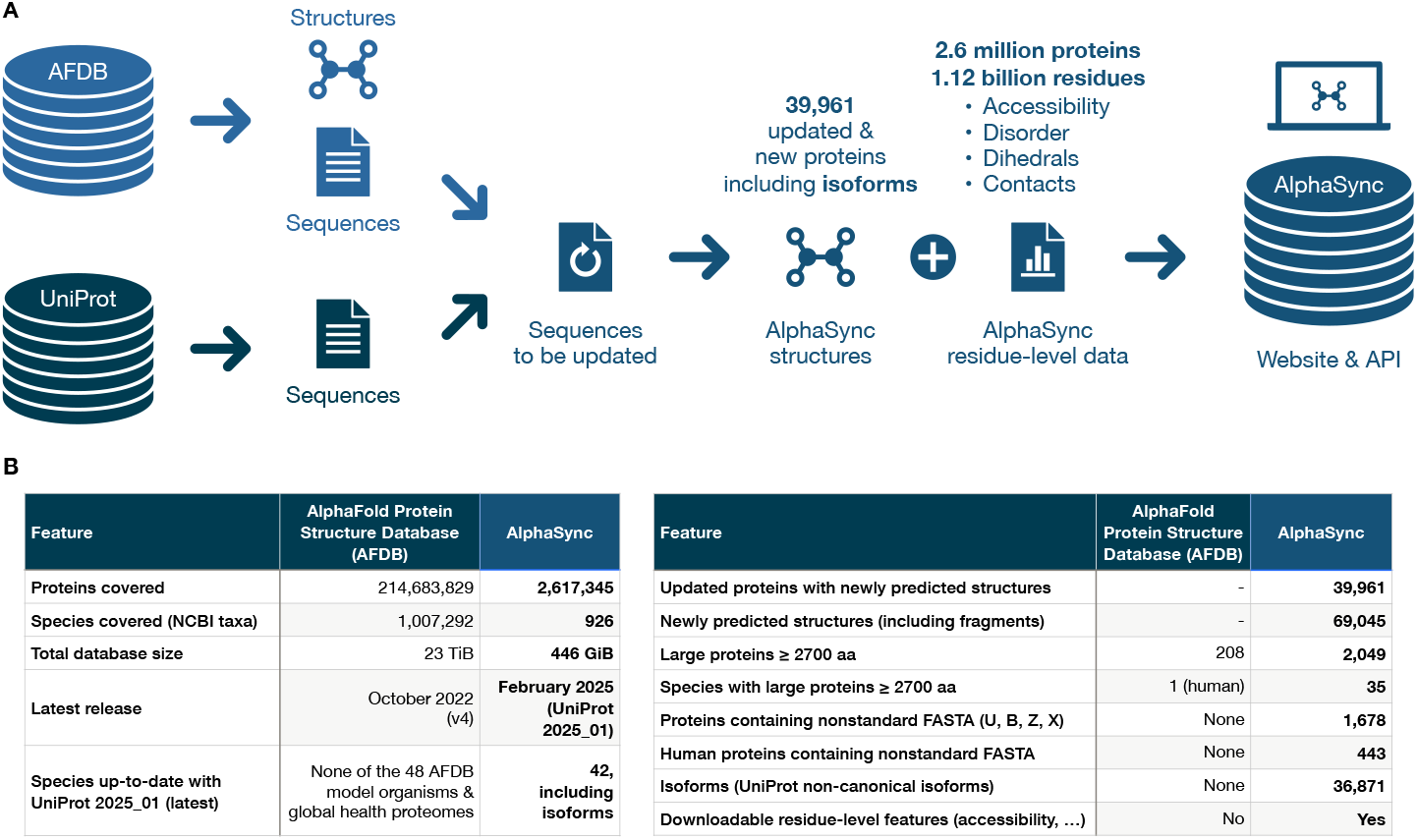
AlphaSync overview. **(A)** Structures are obtained from the AlphaFold Protein Structure database (AFDB) and their sequences extracted. These are then compared to sequences from the latest UniProt release. Any UniProt sequences for which no structure is available in AFDB are selected for AlphaFold-based structure prediction. Once up-to-date structures have been generated for all UniProt proteins for the reviewed and canonical reference proteome proteins for AlphaSync’s species of interest, structural characteristics are computed using various tools, resulting in the detailed quantitative residue-level characteristics and inter-residue non-covalent contacts available via the AlphaSync API and website. **(B)** AlphaSync features that complement the AlphaFold Protein Structure Database. This table shows a comparison of features and database content between the AlphaFold Protein Structure Database (AFDB) and AlphaSync, covering database sizes, species completeness, isoforms, large proteins, special FASTA characters, and release dates.

To achieve complete coverage of reference proteomes, AlphaSync uses several new approaches (**Figure 1B**). Predicting structures for alternative isoforms enables analysis of disease-relevant cases, such as for example vascular endothelial growth factor splice variant VEGF165B (P15692-8), which binds to the KDR receptor but does not activate downstream signaling pathways, does not activate angiogenesis, and inhibits tumor growth^3,7^. Availability of isoform structures may also aid in the assessment of differences for protein and inhibitor design. For large proteins of 2,700 amino acids or more such as BRCA2 and laminin subunit alpha-3, we used the fragmentation approach pioneered for the human proteome by Tunyasuvunakool *et al*.^8^. The human proteome remains the only proteome where structures for large proteins are downloadable from the AlphaFold Protein Structure Database, though these are not available through its web interface. AlphaSync handles large proteins (including titin at 34,350 amino acids) seamlessly and processes fragments into a single representation (see **Online Methods**). Though they should be interpreted with caution, we also predicted structures for peptides below 16 aa, which are excluded from the AlphaFold Protein Structure Database. Rare non-standard amino acid FASTA characters^9^ such as U (selenocysteine), B (aspartic acid or asparagine), Z (glutamic acid or glutamine) and X (unknown amino acids) are replaced with standard amino acids as described in the **Online Methods**. We considered such replacement reasonable, since AlphaFold 2 is notably robust to point mutations^10,11^, and preferable over having no structure predictions for such proteins. The procedure enabled structure prediction for 1,678 additional proteins, 443 of them human. We also noted a median absolute length difference between human canonical and alternative isoforms of 67 amino acids, which unlike smaller mutations should result in a notably different structural model^12^. Finally, the AlphaFold 2 pipeline was split into a parallelizable CPU-only multiple sequence alignment (MSA) generation component and a GPU-only inference part, increasing GPU usage efficiency around fourfold. Despite this increase in efficiency, AlphaSync’s 69,045 structure predictions alone required over 13 years of sequential multi-core CPU and GPU compute time. We hope making these structural models widely available will reduce the need for other researchers to make predictions and decrease overall environmental impact.

While a 3D view of protein structures is invaluable, many research questions require detailed information on individual residues’ properties, their interactions, and their roles within a protein. For example, features such as solvent accessible surface area can be used for antibody design, as epitopes must be exposed to be targetable. Conservation of non-covalent contacts across species, even when structurally equivalent residues are not identical, can help uncover determinants of the protein fold^13^. To address this need, AlphaSync enhances all structures with precalculated residue-level features such as solvent accessibility, intrinsic disorder predictions, dihedral angles, and atom-level intra-protein contacts between amino acids. AlphaSync provides up to 21 data points for each of 1.12 billion residues across 2.6 million proteins, as well as 5 additional data points for over 4.7 billion residue-residue contacts, specifically atom names, contact type, distance and AlphaFold predicted aligned error (PAE). Such information can guide protein design efforts, enable functional interpretation of structures as well as the effects of disease- and phenotype-associated variants. The structure models and non-covalent contacts for a protein of interest can be analyzed and visualized using tools such as the Protein Contact Atlas^14^.Users can find individual proteins on the web interface (https://alphasync.stjude.org/) by searching for real-world biological protein or gene names such as “p53”, as well as by database identifiers such as “P04637”. Searching by sequence beginning at the N-terminus or using wildcards such as “Ran*” is also possible, the latter bringing up hits such as “RANBP1”. Next, AlphaSync will show a list of matches using a heuristic search method. This method first checks whether the term is a precise database accession or gene symbol, prioritizing reviewed UniProt entries, and gradually uses more tolerant matching including in UniProt functional descriptions until it finds one or more hits. A wildcard symbol (*) can be appended to ensure more tolerant matching.

The Display page shows an individual protein or isoform (**Figure 2**). A description of the protein according to UniProt is given at the top of the page, followed by its sequence and a detailed residue-level table. This table, as well as the protein structure, can be downloaded using the “Download” buttons at its top. Clicking on an individual residue within the sequence will jump to its approximate position in the detailed table. On hover (mouseover), detailed information for an individual residue is shown as a tooltip. This includes its AlphaFold local prediction confidence (pLDDT, predicted Local Distance Difference Test), its solvent accessibility, its classification (surface or core, intrinsically disordered or structured), its secondary structure class and dihedral angles such as phi/psi (φ/ψ), as well as the non-covalent contacts this residue makes. The pLDDT confidence score is color-coded as in all AlphaFold publications, with pLDDT ≥ 70 indicating a light blue “confident” prediction and pLDDT ≥ 90 indicating “very high” confidence in dark blue. Similarly, the contacts are color-coded according to the rounded AlphaFold predicted aligned error (PAE) value between the two residues, with a PAE value < 1.5 Å being considered of “very high” confidence and a PAE value ≥ 1.5 Å and < 2.5 Å being considered “confident”. Just as for pLDDT values, contacts of “low” or “very low” confidence should likely not be interpreted^1^. The detailed table provides additional explanation when hovering over individual values, as do the information markers at the top of each column.

**Figure 2:**
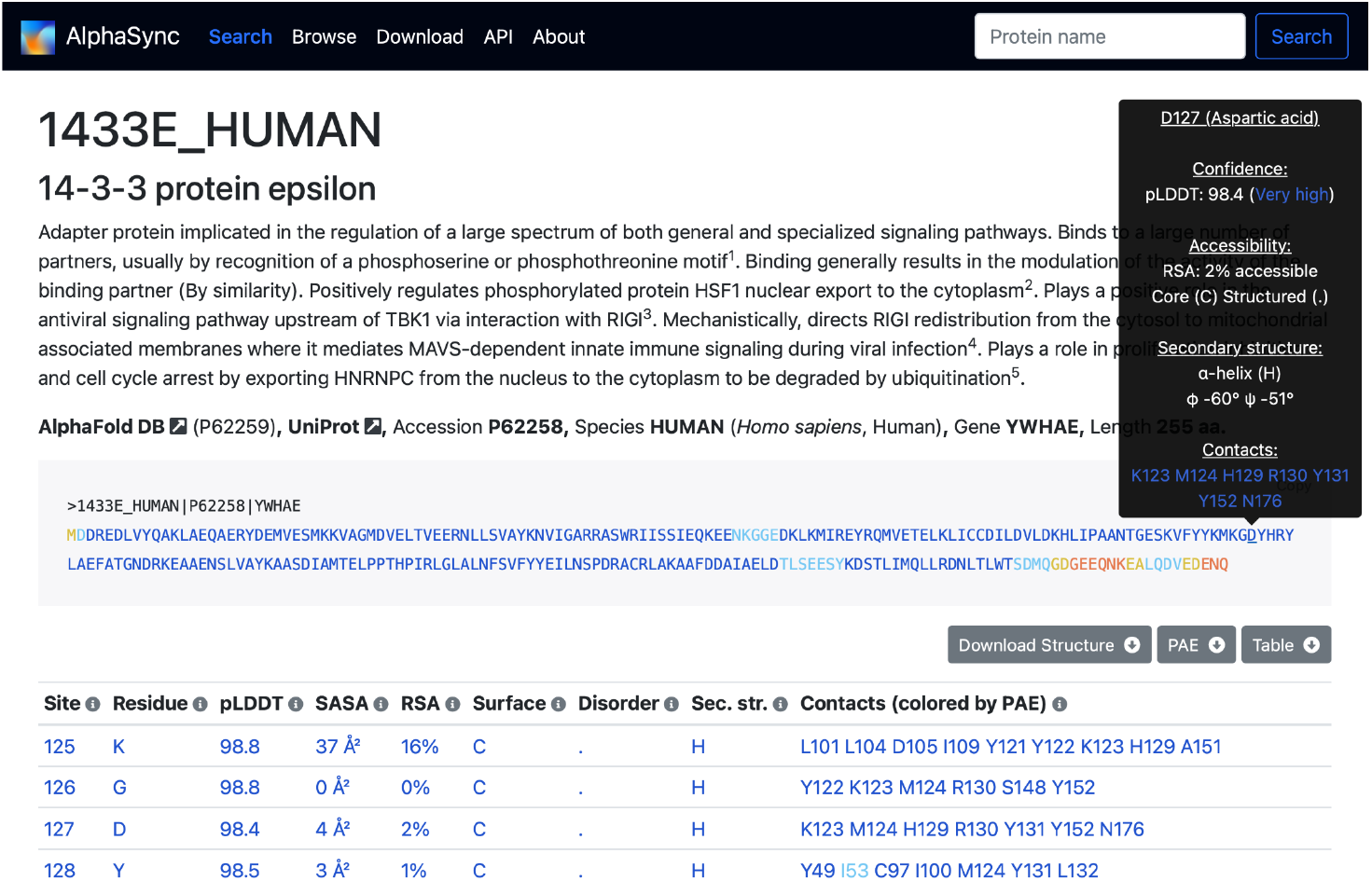
Viewing a protein in AlphaSync. The full protein name and UniProt functional description are shown. When hovering over an amino acid within its sequence, detailed residue-level information and contacts are displayed. A detailed table of residues and their characteristics follows below. For ease of interpretation, explanatory tooltips are provided on hover for all features.

The Download page offers three archives for download. The most important of these contains the 69,045 mmCIF-format structures predicted by AlphaSync, covering a total of 39,961 proteins (large proteins ≥ 2,700 aa were fragmented as detailed in the **Online Methods**). As in the AlphaFold Protein Structure Database, these structures are individually gzip-compressed within a TAR archive and also follow its file naming pattern, allowing direct integration of this archive into existing computational pipelines. Other archives provide predicted aligned error (PAE) score matrices, and the parameters and database versions used to run AlphaFold 2. The Browse page gives an overview of the protein structures and species available in AlphaSync. The species selection dropdown menu at its top lists all 42 completely up-to-date reviewed and canonical reference proteomes (including isoforms) available in AlphaSync, as well as an “all species” option. The API page describes programmatic access to AlphaSync’s database using its REST API. This allows advanced users and developers to easily integrate AlphaSync data within their own analyses, automated workflows and software. The AlphaSync pipeline’s source code can also be adapted for e.g. protein design applications that would benefit from residue-level and contact data. Finally, the About page provides a detailed explanation of AlphaSync, what it provides, its aims and the methods used.

We expect to update AlphaSync with each new UniProt release, currently every two months, as well as to add additional species. We estimate each update to take around 3–7 days, depending on the number of new structures that need to be predicted. We believe our highly approachable, yet powerful resource will provide an essential service to the protein and biomedical research communities, as it can unlock new avenues for protein research and design, particularly at the residue and contact level and at scale.

## CODE AVAILABILITY

AlphaSync’s source code is available at https://github.com/langbnj/alphasync under a BSD-3-Clause licence. The website code is available upon request.

## DATA AVAILABILITY

Newly predicted updated structures for the latest UniProt release can be downloaded at https://alphasync.stjude.org/download under a CC BY 4.0 licence. Also available for download are predicted aligned error (PAE) scores and the exact AlphaFold parameters used to predict structures. AlphaSync’s residue-level data can be programmatically retrieved via the AlphaSync API at https://alphasync.stjude.org/api.

## ACKNOWLEDGEMENTS

We thank Inês Chen and Duccio Malinverni for their helpful input, Besian I. Sejdiu for providing early access to the Lahuta software (https://bisejdiu.github.io/lahuta/) and Chad Burdyshaw for help deploying AlphaFold. We also thank other members of the Babu group for helpful input and discussions and acknowledge ALSAC for financial support.

## FUNDING

AlphaSync’s development is supported by ALSAC.

### CONFLICTS OF INTEREST

None.

## SUPPLEMENTARY TABLE LEGENDS

**Supplementary Table 1: Species included in AlphaSync and their degree of completion**. This table shows the 926 species with structures available in AlphaSync, as well as the number of structures available and those still missing from its complete reviewed and canonical reference proteome. It also clearly marks the 48 model organisms and global health proteomes selected in the AlphaFold Protein Structure Database that we prioritized, as well as the 42 fully completed species (including isoforms) in AlphaSync.

**Supplementary Table 2: AlphaFold 2 parameters used for structure prediction**. This table shows an overview of the parameters, software and database versions used for AlphaFold 2 structure prediction in AlphaSync.

## ONLINE METHODS

### Overview

As described in the main text, AlphaSync begins with structure predictions from the AlphaFold Protein Structure Database (AFDB)^1^, extracts their sequences and identifies sequences that have since been updated in the latest UniProt release (and for which no structure model is yet available). It then predicts new structures for these novel sequences. It also enhances all structures for a set of 2.6 million proteins from species of high interest by calculating residue-level characteristics and predicting intra-protein non-covalent contacts.

### Species and proteins included

AlphaSync begins with a subset of the AlphaFold Protein Structure database, which is extraordinarily large at over 214 million sequences and around 23 TiB of data. Due to resource constraints, we instead focused on a subset of species which we considered of importance to a large number of researchers, covering 2,617,345 proteins from 926 species, 42 of which are covered completely, including isoforms.

We began by focusing on the 48 species included in the AlphaFold Protein Structure Database’s list of model organisms and major human pathogens (“Global Health Proteomes”). From this list, we selected all species for which fewer than 1,000 new structures were needed to arrive at a completely up-to-date canonical reference proteome (one sequence per gene), plus all its reviewed proteins. This resulted in a set of 42 out of 48 species for which we were able to achieve complete up-to-date reviewed and canonical reference proteome coverage in UniProt 2025_01, including UniProt isoforms.

Additionally, we included many proteins from the 200 species that make up the Ensembl Comparative Genomics pipeline (Ensembl Compara release 108, October 2022)^2^. These species are mostly vertebrates and of low-to-medium evolutionary distance to humans. For these species, any protein that had a high-confidence human ortholog, either one-to-one or its best-matching paralog in one-to-many orthology situations (https://useast.ensembl.org/info/genome/compara/homology_types.html), and whose sequence had an exact match in the AlphaFold Protein Structure Database, was included in AlphaSync for calculation of residue-level features and contacts. All high-confidence one-to-one orthologs to human proteins were included, as well as a single “best” one-to-many ortholog for each species that was chosen based on whole-genome alignment (https://www.ensembl.org/info/genome/compara/Ortholog_qc_manual.html), synteny and sequence identity (https://www.ensembl.org/info/genome/compara/homology_method.html). The exact sorting criteria were, in descending order of importance: 1. Ensembl Compara’s high-confidence orthology classification (yes/no), 2. the average of the ortholog’s gene order conservation and whole genome alignment scores, 3. its gene order score, 4. its whole genome alignment score, 5. percent sequence identity for the query sequence, and 6. percent sequence identity for the target sequence. Using these criteria, the highest-ranked ortholog was then selected as the single “best” one-to-many ortholog. This ensures that each of the 200 representative eukaryotic species included in the Ensembl Comparative Genomics pipeline is represented by at most a single protein in a given homology cluster and keeps the size of the AlphaSync database manageable. Except for those species included in the list of 48, no new structures were predicted for the 200 Ensembl Compara species, and they are therefore not completely covered or up-to-date.

We also included all reviewed proteins from UniProt (i.e. UniProtKB/Swiss-Prot) for whose sequences there was a structure available in the AlphaFold Protein Structure Database in all above species, leading to matches across 926 species, and calculated residue-level properties and contacts for them. However, no new structures were predicted for this set, and it is therefore not completely covered or up-to-date.

We next added isoforms of reviewed and canonical reference proteome proteins, beginning with structures for human isoforms. While these had previously been comprehensively predicted by Sommer *et al*.^3^ in December 2022, we found that this dataset covered only 65.1% of current isoform sequences for human reviewed and canonical reference proteome proteins in UniProt release 2025_01. This dataset also used performance-optimizing approaches such as early stopping when one of the five AlphaFold 2 models reported an average pLDDT below 70, resulting in no structure prediction. A second dataset based on UniProt release 2022_03 likewise had a coverage of only 67.4% of up-to-date human isoforms^4^. For consistency, we decided to produce new structure predictions using the standard AlphaFold 2 pipeline, including its full five-model ensemble approach from which the best structure is selected.

### Synchronizing UniProt with the AlphaFold Protein Structure Database

AlphaSync’s aim is to provide protein structures whose sequences are synchronized with the latest UniProt release (currently 2025_01). To achieve this, we first obtain the latest UniProt protein sequences and annotation for the species and proteins mentioned above using UniProt REST API calls while also retrieving isoforms (e.g. “https://rest.uniprot.org/uniprotkb/stream?format=json&includeIsoform=true&query=organism_id:9606” for human). We separately retrieve all reviewed UniProt proteins using the REST API. We further obtain canonical reference proteomes from the UniProt FTP server (e.g. “https://ftp.uniprot.org/pub/databases/uniprot/current_release/knowledgebase/reference_proteomes/Eukaryota/UP000005640/UP000005640_9606.fasta.gz” for human). The final set of proteins for the 48 model organism and global health proteome proteins that we aim to cover completely therefore consists of both reviewed and canonical reference proteome proteins.

An interesting aspect of the AlphaFold Protein Structure Database is that it contains one structure per UniProt accession, regardless of whether its sequence is unique. As a result, the average number of structures we observed per unique accession within our subset of species and proteins was 2.6. While it is tempting to speculate that these might at times include alternative conformations, AlphaFold and related methods tend to be severely limited in their ability to predict more than one structural state, even for the small number of proteins that are known to adopt multiple conformations^5^. To arrive at a single structure per sequence, we selected the structure with the highest average pLDDT score, and in the very rare case of pLDDT ties, we selected the structure with the alphanumerically lowest protein accession.

For each protein, we test whether one or more structure predictions are already available in the AlphaFold Protein Structure Database for its sequence. Since the only input to AlphaFold 2 is a protein sequence, this step can retrieve proteins with unexpected accessions and even from other species. This allows us to make full use of the predictions within the AlphaFold Protein Structure Database. If a structure is available, we retrieve it and calculate residue-level features and non-covalent contacts for it as described below. If not, we first run a new structure prediction using AlphaFold 2 as described below.

### Predicting new structures using AlphaFold 2

We used the most recent AlphaFold 2 version available, 2.3.2, as well as the most recent sequence databases available (see below for versions). We opted not to use AlphaFold 3 for consistency with the AlphaFold Protein Structure Database, and since its prediction quality improvement for monomeric structures is very slight^6^, as its development was primarily focused on predicting multimeric protein complexes. AlphaFold 3 also requires more care in terms of providing the correct ions, ligands and modified residues, as it is capable of explicitly modeling these. AlphaFold 2 therefore seemed the most robust choice. Additionally, while the terms of use for AlphaFold 3 structures are strongly limited (https://alphafoldserver.com/output-terms), AlphaFold 2’s permissive licensing allows full use of the structures in AlphaSync including for molecular docking and commercial applications.

We split the AlphaFold 2 pipeline into two parts: the easily parallelizable (multi-core) CPU-only multiple sequence alignment (MSA) generation component, and the GPU-only inference component. This split greatly increased GPU usage efficiency by a factor of around 4. As recently done by the ColabFold project^7^, we opted to use PDB100^8,9^ rather than PDB70^10^, greatly expanding the set of possible template structures and improving the template search due to the use of FoldSeek and AFDB itself in the construction of PDB100. One reason for using PDB70 in the original AlphaFold 2 seems to have been to avoid overfitting to template structures during training and benchmarking, an incentive which applies much less strongly in inference use^11^. For a random test set of 339 human single-fragment proteins (< 2,700 aa), we observed slightly better average pLDDT values for structures predicted with updated input datasets, rising from a median of 76.4 across structures to 76.6, indicating higher prediction confidence. For a different random test set of 1,725 human structures, changing from PDB70 to PDB100 while using updated input datasets further led to a rise from a median of 74.1 to 74.3. Likewise, for a set of 6,647 structures from human, mouse, *D. melanogaster, C. elegans* and *S. cerevisiae*, we observed an increase in average pLDDT from a median of 73.4 to 73.6. We also used the latest PDB structures available, with a “max_template_date” of 2024-11-25 and the latest structure being from 2024-11-15.

For structure prediction, we first ran AlphaFold 2.3.2 with either 12 (“high_cpus”) or 4 (“default”) CPU cores and a limit of 180 GB of RAM per job for the CPU-only multiple sequence alignment (MSA) generation step that we split off from AlphaFold 2’s standard pipeline, using mainly HPE ProLiant XL225n Gen10 Plus and XL170r Gen9 nodes (AMD EPYC 7542 and Intel Xeon E5-2680 CPUs, respectively). We then ran the second GPU-based inference step using four CPU cores, a limit of 120 GB of RAM and a single GPU on NVIDIA DGX A100 nodes.

We begin multiple sequence alignment (MSA) generation for structure prediction with the following software and source database versions and parameters: hhblits 3.3.0 (default), uniref30 2023_02, full BFD (bfd_metaclust_clu_complete_id30_c90_final_seq.sorted_opt), uniref90 2024_05, mgnify 2024_04, and pdb100 pdb100_foldseek_230517. In a fraction of cases (566 out of 69,045, or 0.8%, e.g. for Q922H1) the above parameter combination consistently failed, as did AlphaFold 2.3.2’s default combination of older databases and parameters (e.g. for Q86VQ6). These failures occurred at hhblits’s merging step between BFD and uniref30 results. We noted that DeepMind had used various combinations of database versions in the published AlphaFold 2, human proteome and AlphaFold 3 articles^6,12,13^, as well as consistently using a specific beta version of hhblits (3.0-beta.3 from 2017-07-14), which uses hardcoded parameters lower than those used by AlphaFold 2.3.2’s hhblits 3.3.0 (“maxseq” 65,535 rather than 1 million, for instance). The human proteome paper did not use any version of uniref30 (also referred to as uniclust30). Presumably these changes were necessary for successful proteome-wide structure prediction.

We ultimately implemented a consistent sequential downgrading approach which, on structure prediction failure, first attempts to lower the “maxseq” parameter given to hhblits 3.3.0 to that hardcoded in hhblits 3.0-beta.3 (from 1 million to 65,535, “low_limits”). Where this fails, we next attempt to run hhblits 3.0-beta.3 with default parameters, followed by “low_limits”. If all hhblits approaches were unsuccessful, we next downgrade the version of uniref30 used from 2023_02 to 2022_02, 2021_03, 2020_06 and subsequently 2018_08, again trying each hhblits approach for each uniref30 version. In practice, uniref30 2021_03 was the oldest version necessary for successful prediction (for I6XD69 fragment 5 only, residues 801–2200). The final, successful parameters (which are identical for 99.2% of structures) are recorded in a JSON file for each structure and are downloadable from the AlphaSync website as a bulk archive or for individual proteins, and statistics are given in a supplementary table (**Supplementary Table 2**). We found this sequential downgrading approach to be fully robust for all sequences, allowing us to achieve complete proteome coverage including isoforms for the first time for currently 42 of the 48 AFDB model organisms and global health proteomes we aimed for.

### Protein fragmentation and tolerance of non-standard FASTA characters (U/B/Z/X)

To achieve complete coverage of proteomes, AlphaSync uses several approaches (**Figure 1B**). For large proteins of 2,700 amino acids or more, we used the fragmentation approach pioneered for the human proteome by Tunyasuvunakool *et al*.^13^. This approach splits a large protein into fragments of 1,400 aa beginning at the N-terminus and steps forward by 200 aa until the C-terminus is reached, with the final fragment being shortened if needed. The human proteome remains the only proteome where structures for large proteins are downloadable from the AlphaFold Protein Structure Database, though these are not available via its web interface. AlphaSync processes fragments into a single representation by averaging properties such as solvent-accessible surface area (SASA), relative accessible surface area (RSA) and AlphaFold’s pLDDT confidence score, while discarding all residues within 200 aa of artificial termini introduced by fragmentation. For secondary structure classification, the majority class is used, with the residue’s pLDDT score (see below) used as a tiebreaker and the more confident residue’s class being used. For non-covalent contacts, the union of contacts is formed across fragments (while discarding residues within 200 aa of artificial termini) and their distances averaged.

Rather than not providing any structure prediction for proteins that contain them, rare non-standard amino acid FASTA characters^14^ such as U (selenocysteine), B (aspartic acid or asparagine), Z (glutamic acid or glutamine) and X (unknown amino acid) are replaced with standard amino acids. As noted in the main text, we considered this unproblematic since AlphaFold 2 has been noted to be highly robust to point mutations. Selenocysteine (U) was replaced with cysteine (C). The ambiguous characters B and Z were replaced with asparagine (N) and glutamine (Q), respectively, with the reasoning that these polar amino acids are less likely to have a drastic effect on protein structure than the charged alternatives, aspartic acid (D) and glutamic acid (E).

Additionally, we found that the majority of non-covalent contacts made by D and E within AlphaSync were polar rather than ionic, making it likely that these polar contacts could be maintained by N and Q (data not shown). Finally, unknown amino acids (X) were replaced depending on their sequence context. Terminal individual occurrences or repeated stretches of X were replaced with flexible glycine linker sequences (GGGGS repeated to the same length) to maintain flexibility and solubility^15^. However, internal stretches of up to three X were replaced with the same number of alanine (A) to maintain potential secondary structures. These choices should ensure maintenance of overall structure in these regions. Taken together, these methods allowed us to achieve full reviewed and canonical reference proteome coverage including isoforms for 42 species, including human, mouse and other model organisms.

### Residue-level features

The residue-level features are calculated using several algorithms. First, we determine secondary structure classes as well as absolute solvent-accessible surface areas in Å^2^ (SASA) using DSSP version 4.2.2.1^16,17^. These absolute accessibility values are then converted into relative surface accessibility (RSA) values using reference values for amino acid surface areas within a polypeptide chain from Tien *et al*.^18^. Residues that are ≤25% exposed according to their relative solvent-accessible surface area (RSA) are considered *core* or *buried* (used interchangeably), while residues with RSA >25% are considered *surface* residues^19^. Residues with an average RSA across a ±10 residue window (RSA_10_) of ≥55% are considered to be part of an intrinsically *disordered* region, while residues with RSA_10_ <55% are considered *structured*^20^. Any RSA values above 1, as might occur at the termini, are displayed as 100% on the AlphaSync website.

Next, comprehensive dihedral angles, including side chain chi (𝒳) angles, are calculated using the BioPython Bio.PDB package^21^. Proline isomerization states are determined as *trans* (common) or *cis* (rare) based on their omega angles, with absolute values ≤ 50º classed as *cis* and absolute values ≥ 130º classed as *trans*. Values in between are considered indeterminate (blank). Finally, intra-protein non-covalent atomic contacts are detected using the Lahuta Python package with default values (https://bisejdiu.github.io/lahuta/). Lahuta reports intra-protein atom-level contacts between residues based on atomic distances, including the type of contact (e.g. “IonicContacts”). The structure models and non-covalent contacts for a protein of interest can then be analyzed and visualized using tools such as the Protein Contact Atlas^22^.

### Numbers on outdated sequences

For presentation in the main text, we set out to quantify how many human reviewed and canonical reference proteome sequences had been updated between the latest release of the AlphaFold Protein Structure Database (version 4, October 2022) and UniProt 2025_01 (February 2025). While the latest AlphaFold Protein Structure Database version (v4) was released in October 2022, it makes use of older UniProt sequences due to the amount of time involved in calculating structure predictions. To quantify the number of updated sequences, we started from the human reviewed and canonical reference proteome accessions from UniProt 2025_01, excepting small peptides below 16 aa and sequences containing ambiguous or special FASTA characters such as B, Z, U or X, for a total of 20,619 accessions (20,541 unique sequences). For each protein accession, we compared its sequence to that found in the structure file for the same accession in the AlphaFold Protein Structure Database, where available. 19,986 accessions had identical sequences (19,920 unique sequences). 141 accessions (136 unique sequences) were completely new in UniProt 2025_01, including e.g. P0DXC1 for SRSP, “Splicing regulatory small protein” (https://www.uniprot.org/uniprotkb/P0DXC1/history). 492 accessions (492 unique UniProt sequences) were present in both versions but had updated sequences, including e.g. P01106 for MYC (https://www.uniprot.org/uniprotkb/P01106/history). The number of new or updated accessions is therefore 633 out of 20,619 proteins, or roughly 3.1% of the human canonical reference proteome. In terms of unique sequences, the numbers are quite similar at 628 out of 20,541 unique sequences (3.1%).

To assess the disease relevance of these proteins, we looked for clinically significant ClinVar^23^ missense variants associated with genetic disorders in these proteins using ClinVar release 2023-07-22 and the Ensembl^2^ Variant Effect Predictor^24^ (VEP) from release 108, as well as for missense mutations in Cancer Gene Census genes (which are considered causally implicated in cancer) using COSMIC^25^ release 99.

For ClinVar, this identified 60 genes whose canonical reference proteome sequence had been updated between AlphaFold Protein Structure Database version 4 and UniProt release 2025_01: COL1A1 (348 unique clinically significant amino acid variants reported, as a ranking proxy for disease relevance), BRCA2 (171), MEN1 (124), KCNJ11 (65), ATRX (60), GABRG2 (36), HEXB (31), TAFAZZIN (27), CLDN16 (24), SLC4A11 (24), RBM20 (22), CHRNA1 (21), BBS2 (21), F13A1 (17), CNGA1 (16), NIPAL4 (15), SPECC1L (14), GDF5 (12), MYC (11), GPIHBP1 (10), COQ2 (10), FREM2 (9), SLC9A6 (7), SFTPA2 (7), DNAH1 (7), CIC (6), DPH1 (6), RPL10 (6), LAMA3 (5), BCORL1 (5), AXIN2 (5), GBF1 (4), AMPD1 (4), AIP (3), AP1B1 (3), SLC30A2 (2), RFX7 (2), OBSCN (2), PLXNA3 (2), OPN1LW (2), IGHM (2), TAP1 (2), ACTL9 (2), FRRS1L (1), ALDH1B1 (1), SEMA6A (1), HIVEP1 (1), XRCC1 (1), SHANK2 (1), POLR3K (1), C2CD6 (1), NTHL1 (1), AURKA (1), TRPC3 (1), NR1D2 (1), LURAP1L (1), SUMO4 (1), TCEAL1 (1), FOCAD (1), and OTOGL (1).

For COSMIC, this approach identified 12 Cancer Gene Census (CGC) genes. From CGC tier 1 (highest confidence), these were: BRCA2 (1,651 unique recurrent amino acid variants reported, recurring in ≥ 3 samples, as a ranking proxy for cancer relevance), ATRX (1,411), CIC (859), BCORL1 (712), COL1A1 (655), MYC (543), AXIN2 (485), MEN1 (346), and RPL10 (75). From CGC tier 2 (candidate), these were MUC4 (3,716), BIRC6 (1,477), and NTHL1 (100).

These findings led us to conclude that maintaining synchronization between structure and sequence databases is indeed of great importance and relevance in biomedical research, as a considerable number of proteins with clear disease relevance have received sequence updates.

### Web interface

AlphaSync will always show a protein’s latest UniProt accession, sequence and details, as found in the latest UniProt release. Internally, as described above, AlphaSync uses perfect sequence matching to identify pre-existing structures within the AlphaFold Protein Structure Database that match a given sequence. For example, OGT isoform accession O15294-2 is mapped by AlphaSync to the pre-existing unreviewed AFDB structure Q548W1. The AlphaSync protein display page for O15294-2 therefore links out to accession Q548W1 in the AlphaFold Protein Structure Database (AFDB).

The Protein Display page of the web interface (e.g. https://alphasync.stjude.org/display/P62258) shows predicted aligned error (PAE) scores in ångströms (Å). As the PAE matrices produced by AlphaFold 2 can be slightly asymmetric, we used the maximum value of PAE(a,b) and PAE(b,a), where a and b are residue numbers and the first residue is the one aligned on (in reference to which the predicted aligned ångström error range is given). This results in a slightly conservative estimate of error, erring on the higher side, though PAE matrices are generally nearly symmetric.

For non-canonical isoforms of proteins, the isoform’s functional description is shown on the Protein Display page if available, along with a note mentioning “Canonical isoform description”. If no isoform description is available, the canonical isoform’s description is shown instead.

### Database statistics

The total size of the AlphaSync data at the time of writing this manuscript (synchronized to UniProt release 2025_01) is 446 GiB. Each structure prediction took a median of 67 minutes of CPU wall time using 4 cores, and 33 minutes of single GPU wall time on NVIDIA DGX A100 nodes. For 69,045 structures this equates to around 3,213 days of 4-core CPU time and around 1,582 days of GPU time, or around 13.1 years of combined compute time. We hope that making these structures widely available will reduce the need for other researchers to make these predictions, and therefore decrease the overall environmental impact.

## Notes

https://alphasync.stjude.org

https://github.com/langbnj/alphasync

### Competing Interest Statement

The authors have declared no competing interest.

### Summary of Updates

We have updated AlphaSync with additional structure predictions, completing isoform coverage for Mus musculus (mouse), D. melanogaster (fruit fly), C. elegans, and A. thaliana. The total number of structural models predicted by AlphaSync is now 69,045, covering 39,961 proteins (an increase of 13,164). We also made minor improvements to the text.

